# Human papillomavirus 16 E2 repression of TWIST1 transcription is a potential mediator of HPV16 cancer outcomes

**DOI:** 10.1101/2020.09.25.314484

**Authors:** Christian T Fontan, Dipon Das, Molly L Bristol, Claire D James, Xu Wang, Hannah Lohner, Azeddine Atfi, Iain M. Morgan

## Abstract

Human papillomaviruses are causative agents in around 5% of all cancers, including cervical and oropharyngeal. A feature of HPV cancers is their better clinical outcome compared with non-HPV anatomical counterparts. In turn, the presence of E2 predicts a better clinical outcome in HPV positive cancers; the reason(s) for the better outcome of E2 positive patients is not fully understood. Previously, we demonstrated that HPV16 E2 regulates host gene transcription that is relevant to the HPV16 life cycle in N/Tert-1 cells. One of the genes repressed by E2 and the entire HPV16 genome in N/Tert-1 cells is TWIST1. Here we demonstrate that TWIST1 RNA levels are reduced in HPV positive versus negative head and neck cancer, and that E2 and HPV16 downregulate both TWIST1 RNA and protein in our N/Tert-1 model; E6/E7 cannot repress TWIST1. E2 represses the TWIST1 promoter in transient assays, and is localized to the TWIST1 promoter; E2 also induces repressive epigenetic changes on the TWIST1 promoter. TWIST1 is a master transcriptional regulator of the epithelial to mesenchymal transition (EMT) and a high level of TWIST1 is a prognostic marker indicative of poor cancer outcomes. We demonstrate that TWIST1 target genes are also downregulated in E2 positive N/Tert-1 cells, and that E2 promotes a failure in wound healing, a phenotype of low TWIST1 levels. We propose that the presence of E2 in HPV positive tumors leads to TWIST1 repression, and that this plays a role in the better clinical response of E2 positive HPV tumors.

**Importance:** HPV16 positive cancers have a better clinical outcome that their non-HPV anatomical counterparts. Furthermore, the presence of HPV16 E2 RNA predicts a better outcome for HPV16 positive tumors; the reasons for this are not known. Here we demonstrate that E2 represses expression of the TWIST1 gene; an elevated level of this gene is a marker of poor prognosis for a variety of cancers. We demonstrate that E2 directly binds to the TWIST1 promoter and actively represses transcription. TWIST1 is a master regulator promoting EMT and here we demonstrate that the presence of E2 reduces the ability of N/Tert-1 cells to wound heal. Overall, we propose that the E2 repression of TWIST1 may contribute to the better clinical outcome of E2 positive HPV16 positive tumors.

## Introduction

HPV are causative agents in around 5% of all human cancers (1). HPV16 is the most prevalent high-risk (those that cause cancer) type of HPV, responsible for around 50% of cervical cancers and 90% of HPV positive oropharyngeal cancers (HPV16+OPC). The latter has reached epidemic proportion in the past generation (2-5). The HPV genome is circular and around 8kbp in size.

HPV infect basal epithelial cells and upon nuclear entry a host of cellular factors activate transcription from the viral long control region (LCR) (6). The resultant viral transcript is processed into individual gene RNAs that are then translated. The viral oncoproteins E6 and E7 target several cellular proteins and disrupt their functions, including the tumor suppressors p53 and pRb, respectively (7, 8). Both p53 and pRb are transcription factors, therefore the presence of E6 and E7 in cells results in a disruption of host gene transcription that contributes to the oncogenic properties of HPV. HPV use two proteins to regulate replication of the viral genome. The E2 protein forms homodimers via a carboxyl terminal domain and binds to four 12bp palindromic sequences within the viral LCR. Three of these surround the viral origin of replication (9) and via the amino terminus of the E2 protein, the viral helicase E1 is recruited to the A/T rich viral origin of replication (10). E1 forms a di-hexameric helicase that then replicates the viral genome in association with host polymerases (11-14). Following infection, the virus replicates to around 50 copies per cell to establish the infection. This copy number is maintained as the infected cell migrates through the epithelium before amplifying in the upper layers; the L1/L2 structural proteins are then expressed and the viral genomes encapsulated resulting in viral particles that egress from the upper layers of the epithelium (15).

As well as acting as a replication factor, the E2 protein can regulate transcription. Where E2 sites are present upstream from a heterologous promoter such as herpes simplex virus 1 thymidine kinase (tk) promoter, E2 can activate transcription (16, 17). In addition, overexpression of E2 can repress transcription from the viral LCR (18-20). Given the ability of E2 to act as a transcription factor, the ability of E2 to regulate host gene transcription has been studied. E2 can regulate transcription via AP1 (21-25), nuclear receptors (26), and C/EBP (27). Transient E2 overexpression identified global gene changes induced by E2 (28-30). To gain a greater understanding of E2 regulation of host gene transcription by physiologically tolerated levels of E2, we generated stable cell lines expressing E2 and identified E2 induced host gene expression changes. This was originally done in U2OS cells (31, 32). However, we wished to develop a more physiologically relevant model and to do this we used N/Tert-1 cells, foreskin keratinocytes immortalized by telomerase. We generated N/Tert-1 cell lines stably expressing HPV16 E2. For comparison, we also prepared N/Tert-1 cells that contained the entire HPV16 genome, and have previously demonstrated that N/Tert-1 cells support late stages of the HPV16 life cycle making it an appropriate model for the study of HPV16 (33). RNA-seq analysis demonstrated a significant overlap between genes regulated by E2 and those regulated by the entire HPV16 genome (34). Many innate immune genes were repressed by HPV16 E2 and the entire genome, and these genes are also repressed by E6 and E7 expression. We recently demonstrated that one of these genes, SAMHD1, is a restriction factor for HPV16 as it controls the viral life cycle in the differentiating epithelium (35). However, we wished to identify a gene that was only regulated by E2, not by E6/E7. Our RNA-seq analysis predicted that TWIST1 was transcriptionally repressed by E2 and the entire HPV16 genome in N/Tert-1 cells; in addition, TWIST1 was also downregulated in HPV16 positive versus negative head and neck cancer (34).

TWIST1 is a basic helix-loop-helix transcription factor critical for promoting epithelial to mesenchymal transition (EMT) and embryogenesis (36, 37). EMT is a critical process in cancer as this trait is associated with high grade malignancy and resistance to chemotherapeutic agents (38-41). EMT is an epigenetic process that proceeds independently from DNA mutations (42). In addition, EMT promotes immune escape of cancer cells (43).

The potential repression of TWIST1 expression by HPV16 E2 is intriguing as it could play a role in dictating therapeutic outcomes. Several reports have demonstrated that the expression of E2 in HPV16 positive tumors predicts improved survival and repression of TWIST1 would correlate with this improved survival (44-46). Here we demonstrate that TWIST1 is transcriptionally repressed by E2 in N/Tert-1 cells, and that TWIST1 is downregulated in HPV positive head and neck cancer versus HPV negative. The mechanism of E2 repression is not due to DNA methylation, but involves direct binding of E2 to the TWIST1 promoter, repressing transcription and inducing repressive histone markers. We demonstrate that TWIST1 target genes are also downregulated in E2 expressing cells, and that wound healing is compromised. As EMT is a process that occurs during wound healing, the observed change in wound healing suggests that E2 contributes to EMT suppression in N/Tert-1 cells (47). Two HPV16 positive head and neck cancer cell lines were also studied, one that has episomal viral genomes and therefore expresses E2, and the other with an integrated HPV16 genome that has lost E2 expression. In the episomal genome containing cell line, the presence of E2 resulted in decreased levels of TWIST1 when compared with the integrated non-E2 expressing cell line. Finally, neither E6 nor E7 is able to regulate the expression of TWIST1. Overall, our results support the idea that E2 represses TWIST1 expression during the HPV16 life cycle, and that this downregulation persists into HPV16 positive tumors. TWIST1 repression would promote a better patient outcome, a hallmark of E2 expression (44-46).

## Results

### TWIST1 is transcriptionally repressed by E2, but not E6/E7

Our RNA-seq analysis of N/Tert-1+E2 (expressing HPV16 E2 only) and N/Tert-1+HPV16 (containing the entire HPV16 genome) demonstrated that there was a highly statistically significant overlap between the genes regulated by E2 and the entire HPV16 genome (34). In addition, using our analysis of TCGA data, we observed a significant downregulation of TWIST1 in HPV16 positive versus negative head and neck cancers (HNSCCs) (33, 34). Figure 1A summarizes the expression of TWIST1 in both episomal and integrated HPV16 positive versus HPV16 negative HNSCCs. Because integration disrupts E2 gene expression, integration and episomal groups were determined depending on their E2 expression (46, 48). TWIST1 mRNA expression data were obtained from 528 TCGA tumors using cBio Cancer Genomic Portal (49, 50). We have previously characterized HPV16 status and viral integration in these samples (46). TWIST1 mRNA expression was then compiled and reported by HPV16 status (Figure 1A). We found that TWIST1 mRNA is expressed at statistically significantly lower levels in episomal HPV16 positive tumors compared to those that were either integrated or HPV16 negative. There was no statistical difference in TWIST1 expression between integrated and HPV-negative tumors. Figure 1B summarizes our previous data from our RNA-seq analysis in N/Tert-1 cells. TWIST1 was found to be significantly downregulated in N/Tert1 cells expressing HPV16 E2 or the entire viral episome, compared to parental control cells (33, 34).

**Figure 1.**
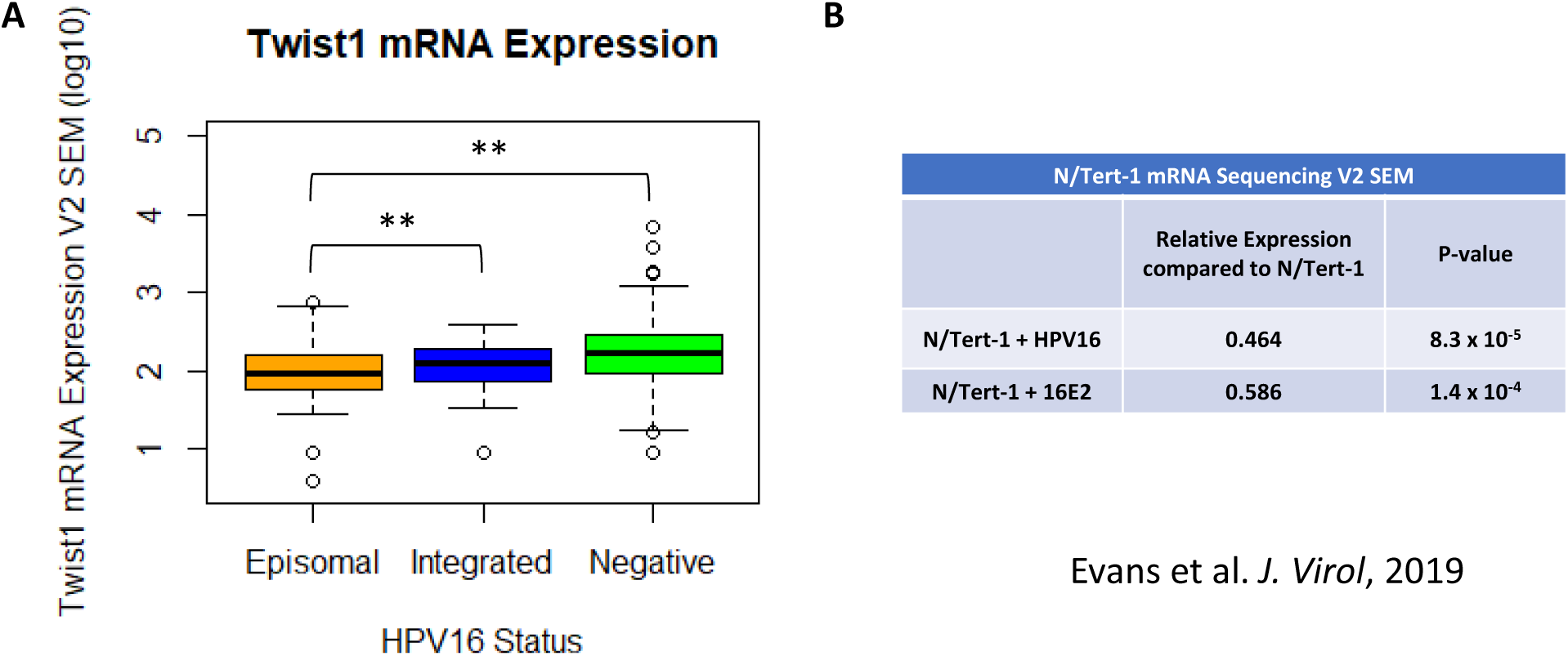
TWIST1 expression is downregulated by HPV16 E2. (A) 528 samples were previously evaluated for HPV genome status utilizing E2-E7 ratio of mRNA-sequencing reads (48, 76). 466 samples were HPV negative. 40 samples retained HPV16 episome status and 17 samples had integrated HPV16 genomes. Firehose Legacy Twist1 mRNA expression data was obtained using cBioPortal. TWIST1 mRNA between HPV16 Status was then compared using two-tailed Student’s T-Test. Vertical axis in log(10). (B) N/Tert-1 RNA-sequencing results adapted from (34). TWIST1 expression was compared to parental N/Tert-1 cells with no HPV16 or HPV16 E2. *P-value <0.05 using Bonferroni correction when applicable.

Figure 2A validates the downregulation of TWIST1 RNA expression in N/Tert-1 cells expressing E2 or the entire HPV16 genome (compare lane 2 and 3, respectively, with lane 1). We have also previously reported the overexpression of HPV16 E6 and E7 in the same cell background, and these oncogenes did not alter the expression of TWIST1 (lane 4). This is in contrast with our prior study of innate immune response gene regulation by HPV16 which demonstrated that E2, E6 and E7 can all repress these genes (34). Therefore, TWIST1 is the first gene we have determined to be likely exclusively regulated by E2 during the HPV16 life cycle. To confirm that the RNA expression was reflected at the functional protein level, western blots were carried out for TWIST1 (Figure 2B). There is a clear downregulation of TWIST1 protein expression in cells expressing E2 or HPV16 (compare lanes 2 and 3, respectively, with lane 1) but not in cells expressing E6/E7 (compare lane 4 with lane 1). This was repeated another two times and the results quantified (Figure 2C); both E2 and HPV16 induce a significant reduction in TWIST1 protein levels, while E6/E7 do not.

**Figure 2.**
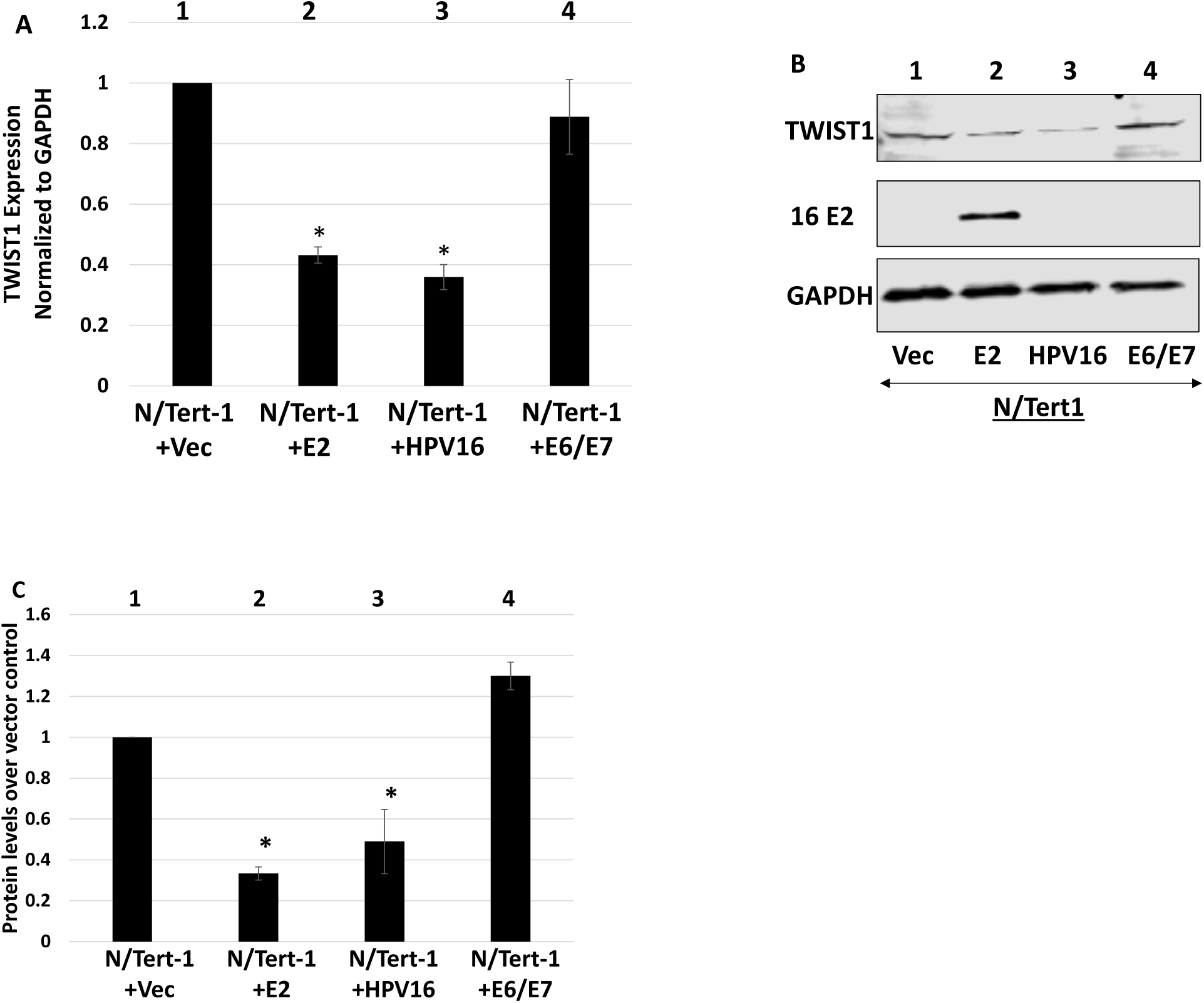
TWIST1 RNA and protein levels are downregulated by E2 and the HPV16 genome in N/Tert-1 cells. (A) qRT-PCR of N/Tert1 lines with E2 (lane 2), HPV16 (lane 3), E6 and E7 (lane 4) or empty pCDNA3.1 vector (lane 1). Results are expressed as fold change from that observed in the vector control N/Tert-1 cells (lane 1). (B) Western blot analysis was carried out on protein extracted from N/Tert-1 cells with empty vector (lane 1), or those with E2, HPV16 or E6 and E7 (Lanes 2-4). GAPDH is shown as an internal control. Western blots were visualized using a Li-Cor system. (C) Western blots were quantitated and TWIST1 protein expression was calculated relative to vector using ImageJ. Data in panels A and C represent the averages of at least 3 independent experiments, and error bars indicate standard error of the mean. *, *P* < 0.05

### E2 binds to the TWIST1 promoter region and directly represses transcription

Our previous work demonstrated that E2 represses the transcription of innate immune genes by regulating the DNA methylation of the corresponding promoters (34). To determine whether E2 is also regulating TWIST1 expression via DNA methylation of the promoter, we treated the cells with 1μM Decitabine, a DNA methylase inhibitor that relieves E2 mediated repression of innate immune genes (Figure 3). Figure 3A demonstrates that the drug has worked as there is an increase in both MX1 and IFIT1 when Decitabine is added, as we observed previously (34). The increase occurs in the vector control cells (compare lanes 3 and 4 versus 1 and 2) but the relief of repression is greater in the presence of E2 (compare lanes 7 and 8 versus 5 and 6) and HPV16 (compare lanes 9 and 10 versus 11 and 12); these results duplicate those we observed previously (34). However, the TWIST1 levels were not altered in any of the cell lines following treatment with Decitabine. Therefore, TWIST1 is regulated by E2 differently from innate immune response genes and is independent from DNA methylation.

**Figure 3.**
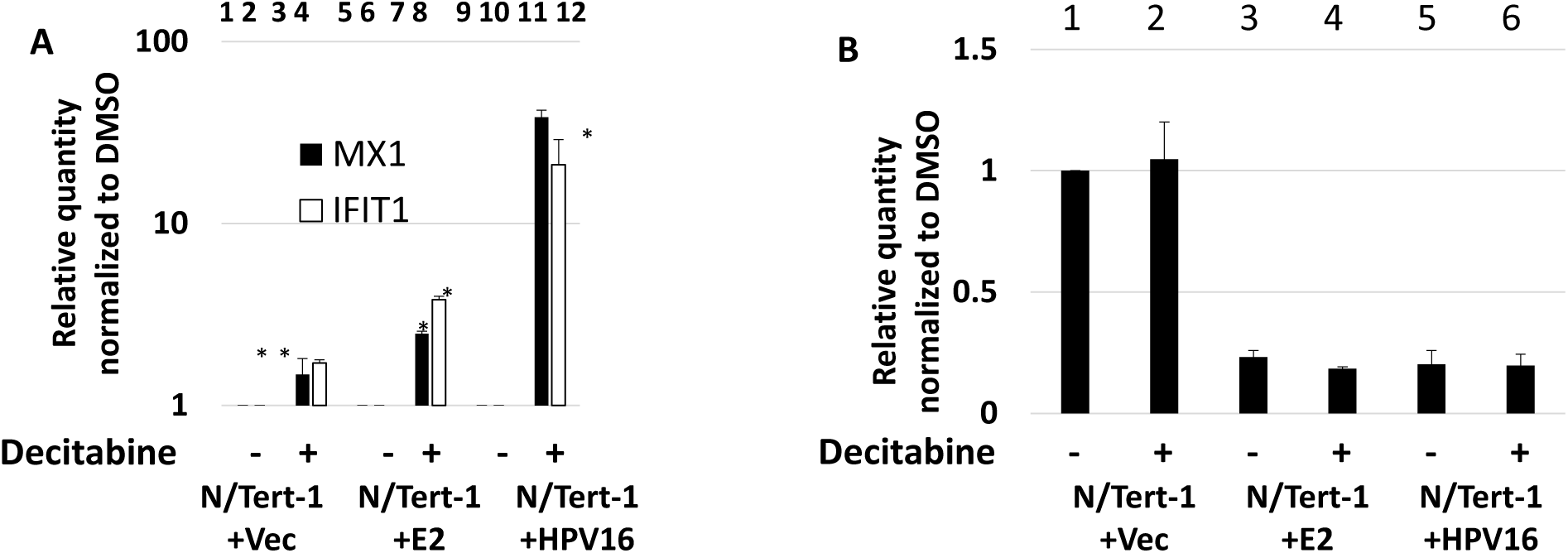
TWIST1 expression is not repressed by E2 via methylation of the gene promoter. N/Tert-1 cells were treated with 1μM Decitabine (5-aza-cytidine) for 72 hours. Afterwards, cells were harvested and processed for RNA which was reverse transcribed into cDNA. (A) qRT-PCR on N/Tert-1 cells for innate immune genes MX1 and IFIT1 illustrates robust restoration of expression following Decitabine treatment. Results are expressed as fold change from that observed in the untreated, vector control N/Tert-1 cells. (B) The same cDNA was then analyzed for TWIST1 expression. Unlike the innate immune genes, Decitabine does not restore TWIST1 repression. Data in panels A and B represent the mean of two independent experiments, and error bars indicate standard error of the mean. *, *P* <0.05

We next investigated whether E2 could directly repress transcription from the TWIST1 promoter. A construct containing the TWIST1 promoter upstream from the luciferase gene (pTWIST-luc) was co-transfected with E2 into N/Tert-1 cells to determine whether E2 can repress expression directly from the TWIST1 promoter (Figure 4A). The expression of E2 resulted in a ∼10-fold reduction in luciferase activity demonstrating that E2 directly represses this promoter. To determine whether E2 can bind directly to the TWIST1 promoter, we carried out a chromatin immunoprecipitation assay (ChIP) using N/Tert-1+Vec, N/Tert-1+E2 and N/Tert-1+HPV16 cells using a sheep E2 antibody as previously described (51, 52) (Figure 4B). There was a significant increase in signal in N/Tert-1+E2 and N/Tert-1+HPV16 when compared with the signal obtained with N/Tert-1+Vec (compare lanes 2 and 3, respectively, with lane 1), demonstrating that E2 binds to the TWIST1 promoter region when overexpressed but also in the context of the entire HPV16 genome. We next investigated whether the levels of repressive chromatin markers implicated in TWIST1 regulation are changed in the presence of E2. H3K9me2 is a repressive chromatin marker involved in the regulation of Twist1 expression (53, 54) and the levels of this marker on the TWIST1 promoter are increased in N/Tert-1+E2 and N/Tert-1+HPV16 when compared with N/Tert-1+Vec (Figure 4C, compare lanes 2 and 3, respectively, with lane 1). These results suggest that E2 interacts with the TWIST1 promoter directly and modifies the local epigenetic environment around the promoter, leading to a reduced level of transcription.

**Figure 4.**
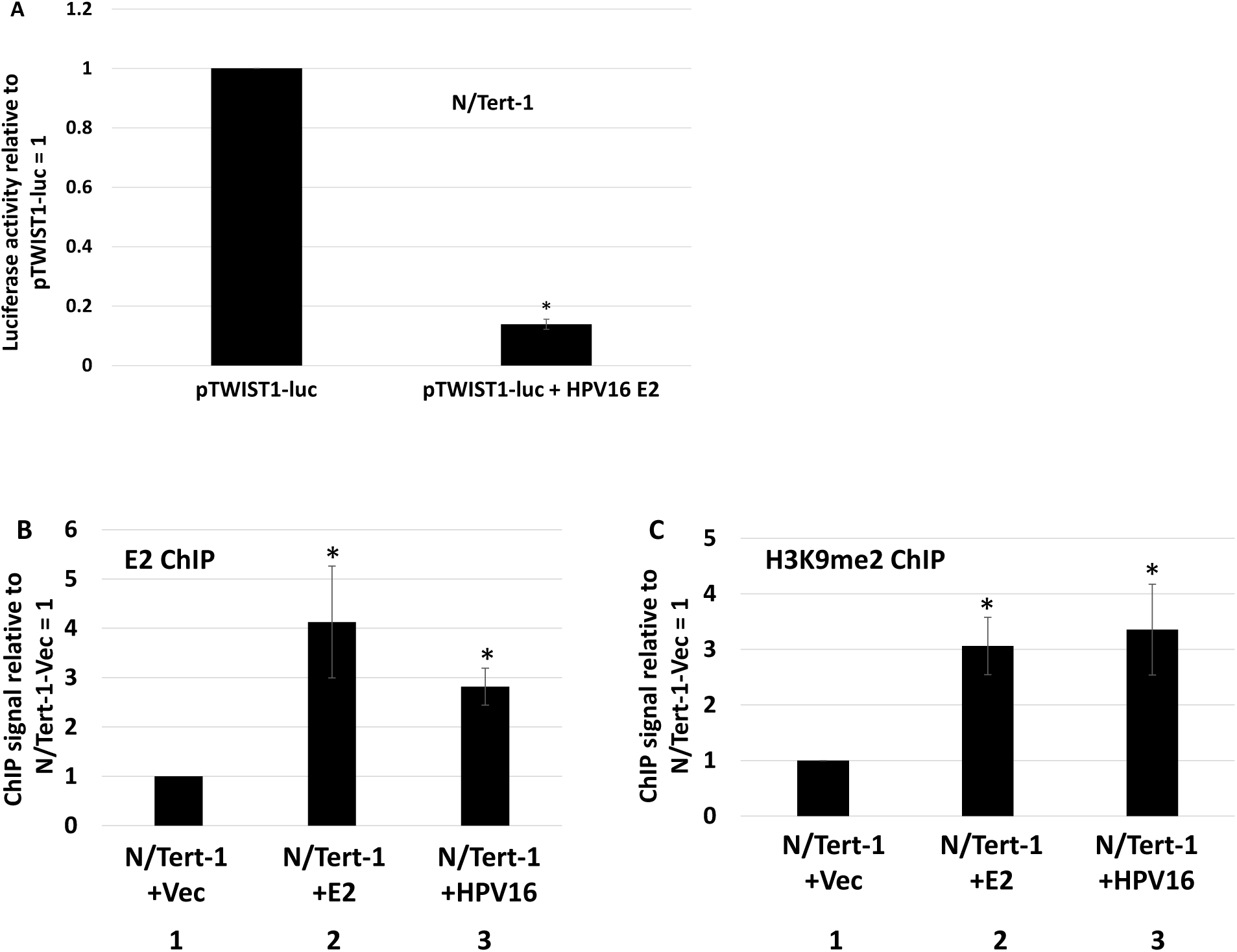
E2 binds to the TWIST1 promoter and actively represses transcription. (A) N/Tert1 cells were transfected with 1 μg of pTWIST1-luc alone or with 1 μg HPV16 E2. Forty-eight hours after transfection, a luciferase-based assay was utilized to monitor levels of TWIST1 promoter activation. Data was obtained as relative fluorescence units (RFU), normalized to total protein concentration as monitored by a standard bovine serum albumin (BSA) assay, and presented as relative to pTWIST1-luc only transfection. (B) Chromatin immunoprecipitation of E2 onto the TWIST1 promoter. In both E2 (lane 2) and HPV16 cells (lane 3), E2 was observed directly binding to the promoter region of TWIST1. (C) E2 binding at the Twist1 promoter leads to accumulation of repressive histone marker H3K9me2. Once again, this repressive marker was upregulated in both E2 (lane 2) and HPV16 expressing (lane 3) N/Tert-1 cells. Results were normalized to input DNA and expressed as fold change over vector control N/Tert-1. Data represent the mean of at least three independent experiments, and error bars depict standard error of the mean. *, *P <* 0.05

### E2 expression reduces the rate of wound healing in N/Tert-1 cells

The repression of TWIST1 expression by E2 suggested a suppression of the EMT phenotype in N/Tert-1 cells. EMT is not a defined status, but is a spectrum of phenotypes that range from epithelial through to fully mesenchymal (47). The N/Tert-1+E2 cells do not look appreciably different to N/Tert-1 control cells, therefore we propose that there is a subtle influence of E2 on the EMT status of these cells. During wound healing there is an EMT transition of the wounded epithelia cells that promotes migration and eventual wound closure (55). Our previous studies on U2OS cells demonstrated that the expression of E2 slowed the ability of these cells to close wounds in monolayer cells (32), therefore we carried out “scratch” assays with our N/Tert-1 cells in order to investigate wound healing. Figure 5A shows images of the results from the wound healing experiment. 20 hours following wound induction, the wound is almost completely healed in N/Tert-1+Vec cells (top 3 panels). However, with N/Tert-1+E2 and N/Tert-1+HPV16 there is a failure to close the wound after 20 hours (second and third from top panels, respectively). To determine whether the oncogenes E6/E7 play a role in regulating wound healing, we also determined wound closure in N/Tert-1+E6/E7 cells (characterized and described in (34)). The presence of the viral oncogenes made no difference to the wound healing (compare the bottom three panels with the top three). Therefore, the failure to close the wound is reflective of TWIST1 levels and E2 expression. This assay was repeated several times and the data quantified (Figure 5B). There is a significant delay in wound healing 12 and 20 hours after the initial “scratch” in both the N/Tert-1+E2 and N/Tert-1+HPV16 cell lines when compared with N/Tert-1+Vec cells. To confirm that TWIST1 function is also downregulated in the presence of E2, we monitored expression of TWIST1 target genes Vimentin and N-Cadherin in N/Tert-1+Vec, N/Tert-1+E2, N/Tert-1+HPV16 and N/Tert-1+E6/E7 cells (Figure 5C). The expression levels of these TWIST1 target genes are reflective of the TWIST1 levels in the cell. This demonstrates that there is a functional loss of TWIST1 in E2 and HPV16 expressing N/Tert-1, and this manifests as alterations in the wound healing phenotype.

**Figure 5.**
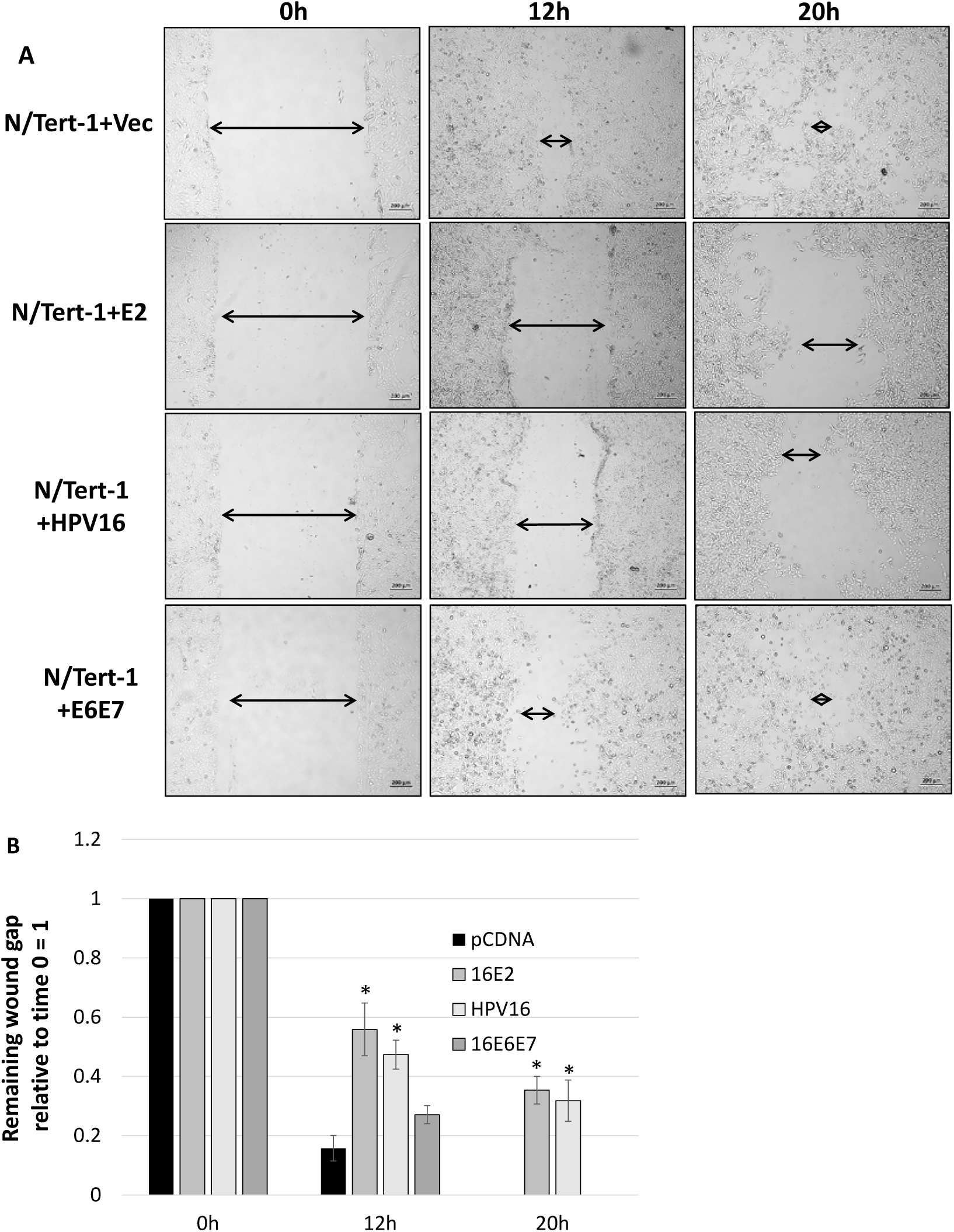
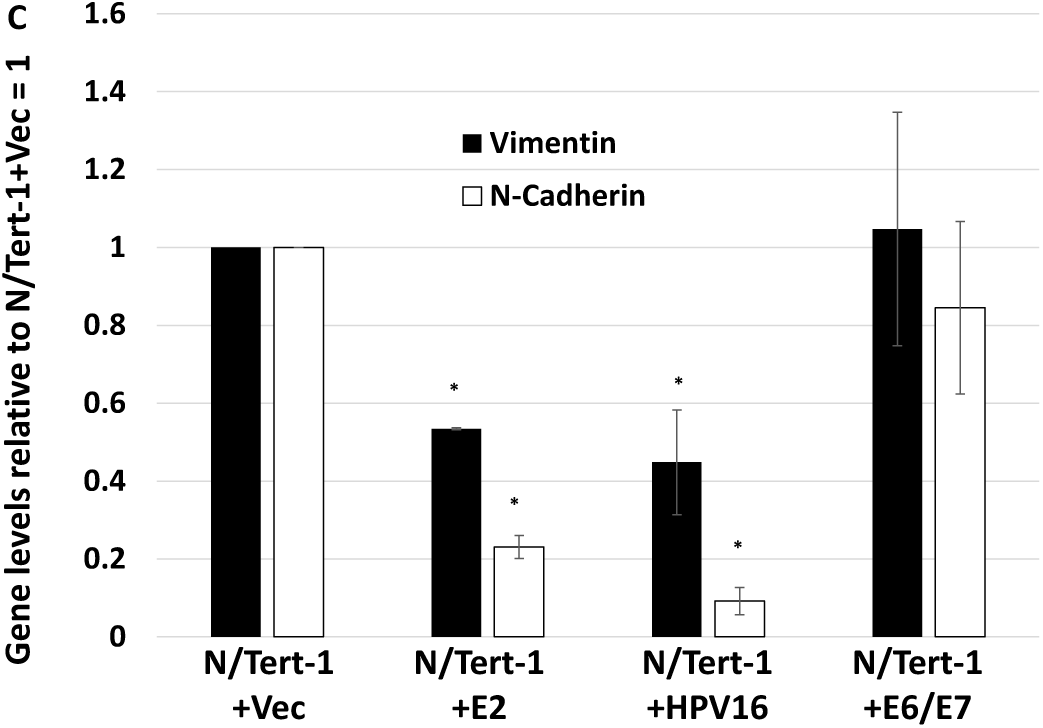
E2 expression correlates with attenuated ability to wound heal. (A) N/Tert-1 cells were plated and allowed to grow to near confluency. Afterwards an ∼1mm scratch was made in the cell monolayer using a pipette tip. The wound gap was imaged and measured at the same field at 0, 12 and 20 hours. Arrows have been added for clarity. Size bars have been added at the bottom-right of each image to illustrate a length of 200 microns. (B) The remaining wound gap was calculated relative to time 0 for each field. By 20h, the Vector and E6E7 cells have completely closed wound gaps while the E2 and HPV16 cells retain considerable gaps. (C) Repression of Twist1 by E2 and HPV16 leads to reduction in EMT marker expression. N/Tert-1 cells were harvested for RNA and processed for cDNA. qRT-PCR was performed for TWIST1 target genes CDH2 and VIM which encode the proteins N-Cadherin and Vimentin respectively. Results are expressed as fold change from that observed in the vector control N/Tert-1 cells (lane 1). Data represent the mean of at least three independent experiments, and error bars depict standard error of the mean. *, *P <* 0.05

### E2 and TWIST1 levels inversely correlate in HPV16 positive head and neck cancer cell lines

Figure 1 demonstrates that there is, on average, less expression of TWIST1 in HPV positive head and neck cancers when compared with HPV negative head and neck cancers. Our results suggest that the expression of E2 is influential in the expression of TWIST1 within the HPV positive group. Previous studies suggested that UMSCC104 contained episomal HPV16 genomes (and therefore retained E2 expression), while UMSCC47 had integrated viral genomes (and therefore had lost E2 expression). We confirmed that UMSCC104 expressed E2 and that UMSCC47 did not (Figure 6A). We next investigated the TWIST1 levels in these cell lines. TWIST1 RNA expression was significantly downregulated in UMSCC104 when compared with UMSCC47 (Figure 6B), and the levels of RNA expression were reflected in a downregulation of TWIST1 protein levels (Figure 6C). The western blots were repeated several times and quantitated; there is significantly less TWIST1 protein expressed in UMSCC47 when compared with UMSCC104. The results in these two HPV16 positive head and neck cancer cell lines support the model of E2 repressing TWIST1 gene expression.

**Figure 6.**
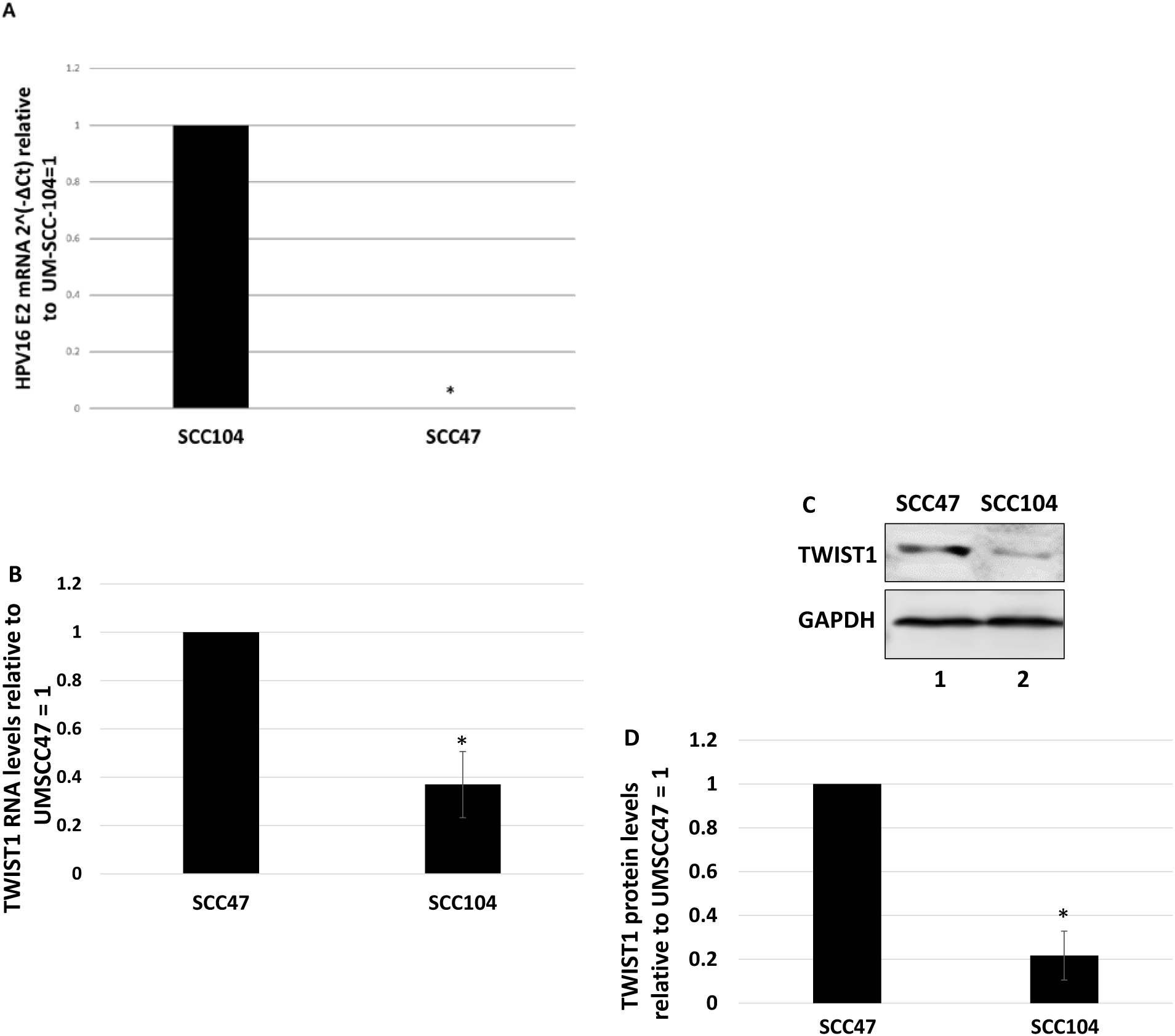
Expression of TWIST1 inversely correlates with E2 expression in head and neck cancer cell lines. (A) qRT-PCR for HPV16 E2 expression in UM-SCC-104 and UM-SCC-47 cell lines. Results are expressed as 2^(-ΔCt)^ using GAPDH as internal control. Integrated UM-SCC-47 exhibited no Ct change over negative experimental control, illustrating no presence of E2 mRNA. (B) The same cDNA in panel A were analyzed for TWIST1 expression. Episomal UM-SCC-104 have lower TWIST1 transcription levels compared to UM-SCC-47. Results are expressed as relative fold change from that observed in UM-SCC-47 cells. (C) Western blot analysis was carried out on protein extracted from the cells in panels A and B. Decreased TWIST1 transcription leads to reduced expression on the protein level in UM-SCC-104 compared to UM-SCC-47 cells. GAPDH is shown as internal control. Western blots were repeated and quantitated and TWIST1 protein expression was calculated relative to UM-SCC-47 protein levels using ImageJ. Data represent the averages of at least 3 independent experiments, and error bars indicate standard error of the mean. *, *P* < 0.05

## Discussion

The HPV E2 protein plays multiple crucial roles during the viral life cycle. It is essential for replication of the viral genome, can regulate the segregation of the viral genome into daughter cells, and has transcriptional properties with the potential to regulate transcription from the viral and human genomes (9, 56). Regulation of host gene expression by E2, and its importance in the viral life cycle, is relatively understudied compared with its role in viral replication and segregation. Our recent work demonstrates that E2 regulates host gene transcription that is relevant during the viral life cycle (34). That report demonstrated an overlap in the ability of E2 and E6/E7 to repress the expression of innate immune genes. In this current report we extend our observations on the transcriptional regulatory functions of E2. We demonstrate that E2 can repress TWIST1 gene expression and that this repression is via a distinct mechanism compared to innate immune gene repression. E6 and E7 did not affect the levels of TWIST1 RNA and protein, therefore it is likely that TWIST1 expression is regulated by E2 during the HPV16 life cycle. For innate immune genes, methylation of the gene promoters played a significant role in repressing expression, but for TWIST1 DNA methylation plays no role. Rather, E2 binds to the TWIST1 promoter and actively represses transcription from this region. E2 also increases H3K9me2 at the promoter, a hallmark of transcriptionally repressed genes previously implicated in regulation of Twist1 transcription (53). To our knowledge, this is the first time E2 has been demonstrated to directly bind to a host gene promoter and induce repressive epigenetic markers. The repression of TWIST1 by E2 resulted in downregulation of TWIST1 target genes Vimentin and N-Cadherin. During wound healing there is an EMT-like transition that promotes wound closure, and wound healing in E2 expressing N/Tert-1 is compromised when compared to parental cells (55).

Repression of TWIST1 expression by E2 may play a role in HPV cancer outcomes. Overexpression of TWIST1 is associated with poorer overall survival in head and neck cancer (57), while E2 expression is associated with improved outcomes in HPV positive head and neck cancer (44-46). We demonstrate here that TWIST1 is downregulated in HPV16 positive tumors that retain E2 expression when compared with HPV16 tumors that have no E2 expression, or are HPV negative. We therefore highlight this correlation between E2 expression and TWIST1 repression which may contribute to the better clinical outcomes of HPV16 positive tumors that retain E2 expression. We demonstrate, in two HPV16 positive head and neck cancer cell lines (one that is E2 positive, one that is negative), that E2 expression inversely correlates with TWIST1 expression which correlates with our observations in N/Tert-1 cells and TCGA datasets. While these lines are not isogenic, they provide a model for studying the repression of TWIST1 in cancer cell lines that is potentially mediated by expression of the E2 protein. Not all E2 positive head and neck tumors have downregulation of TWIST1 compared with E2 negative tumors as evidenced by TCGA data, but there is a significant trend for downregulation. Not only has elevated TWIST1 been associated with poorer survival in several cancers, including head and neck, but TWIST1 protein attenuates the response to chemotherapeutic drugs which provides a rationale for the worse clinical outcomes in high TWIST1 expression patients (47, 55).

The mechanism of E2 repression of TWIST1 expression remains to be fully elucidated. The TWIST1 promoter is methylated in cancer, although this did not correlate with low TWIST1 RNA or protein levels (58). Methylation does not play a role in the mechanism that E2 uses to repress TWIST1 levels as treatment of cells with Decitabine did not relieve TWIST1 repression, therefore E2 uses multiple mechanisms to regulate host gene transcription. We demonstrate here for the first time that E2 binds to and represses transcription from the TWIST1 promoter and modifies chromatin around the promoter start site by inducing elevated levels of H3K9me2, a repressive marker. The transcription factor SP1 is involved in the basal transcription levels from the TWIST1 promoter, and displacement of this factor from the promoter repressed TWIST1 transcription (59). The E2 protein has been shown to displace SP1 from HPV LCRs resulting in transcriptional repression, and may act similarly on the TWIST1 promoter (60, 61). This active repression may be required to block TWIST1 expression as STAT3 is activated by HPV in cervical cancer cells (62), and active STAT3 is an activator of TWIST1 expression, promoting EMT (63). We demonstrate here that there is an increase in the level of the repressive marker H3K9me2 at the TWIST1 promoter in the presence of E2. The chromatin repressor NURD complex can be recruited to the TWIST1 promoter via DOC1, resulting in transcriptional repression via induction of a nucleosome on the TWIST1 promoter. It is feasible that E2 recruits a repressor complex to the TWIST1 promoter in order to regulate transcriptional repression via nucleosome assembly (64). E2 functionally interacts with BRD4, and this interaction is involved in regulating E2 repression of the viral LCR. Again, such a mechanism may be used by E2 to contribute to the repression of the TWIST1 promoter (65). Future studies will characterize the mechanism(s) that E2 uses to repress the TWIST1 promoter. Enhancing our understanding of this mechanism is important as E2 offers a model system for repressing TWIST1 transcription. For example, if H3K9me2 methylation of the TWIST1 promoter is a major contributor to repression by E2 then induction of this modification via drug treatment could be an opportunity to promote a better response of high TWIST1 expressing tumors to chemotherapeutic agents. In addition, if TWIST1 repression is an important mechanism promoting the HPV16 life cycle in epithelial cells, then reversing this repression offers an opportunity to disrupt this process and alleviate HPV16 infections and disease.

Another question is: why does HPV16 repress the expression of TWIST1? While TWIST1 expression is important in mouse embryogenesis, demonstrating an important role in cellular proliferation and differentiation, there is less known about the effect of TWIST1 on epithelial cell differentiation (37). HPV manipulate epithelial differentiation in order to create an environment that supports viral replication (66). While TWIST1 is clearly a promoter of EMT, the effect of this protein on the epithelial differentiation pathway is less clear and is worthy of future study. Perhaps the repression of TWIST1 plays a role in promoting an environment that allows HPV16 replication in the infected epithelium. TWIST1 and HPV16 E2 interact with the NF-κB pathway and downregulation of TWIST1 may promote the ability of E2 to regulate NF-κB, which could be important for the HPV16 life cycle (67, 68). TWIST1 has been shown to downregulate expression of C/EBPα, a protein that E2 functionally interacts with to regulate transcription (27, 69). Therefore, downregulation of TWIST1 expression may prevent disruption of E2’s ability to regulate host gene transcription via C/EBPα during the viral life cycle. Recently, it has been demonstrated that C/EBPα is a crucial factor for epithelial maintenance and prevents EMT, thus E2 downregulation of TWIST1 may enhance the ability of C/EBPα to carry out this function (70). This would also promote the enhanced epithelial status of the E2 positive cells.

In conclusion, this report demonstrates that HPV16 E2 downregulates the expression of TWIST1. This occurs in cells that contain the entire HPV16 genome, and also occurs in HPV16 positive cancers that are E2 positive. Given the important role that TWIST1 plays in EMT, cancer outcomes, and response to chemotherapeutic agents, a fuller understanding of the interaction of HPV16 with TWIST1 is warranted.

## Materials and Methods

### Differential expression in TCGA

Head and neck squamous cell carcinoma (HNSCC) TWIST1 mRNA expression data were obtained from The Cancer Genome Atlas using cBio Cancer Genomics Portal (49, 50). HNSCC tumor samples (Firehose Legacy) were analyzed for HPV status and viral genome integration as previously described utilizing HPV16 E2:E7 ratio of mRNA sequencing reads (46, 71). Samples without available TWIST1 mRNA expression data were omitted. Data for 528 HNSC samples were available. Of these, 522 were identified with available TWIST1 mRNA expression and HPV status which was correlated and reported using R. Statistical analysis was performed using two-way student’s t-test with Bonferroni correction for two comparisons.

### Cell culture

Low passage N/Tert-1 with stably expressing HPV16, 16E2 and 16E6+E7 were generated as previously described and characterized in previous studies.(31-35). These cells were cultured alongside empty vector, drug selected pcDNATM 3 with 111μg/mL G418 Sulfate (Genticin) (Thermo Fisher Scientific). All N/Tert-1 cells were grown in keratinocyte serum-free medium (K-SFM;Invitrogen) with a 1% (vol/vol) penicillin-streptomycin mixture (Thermo Fisher Scientific) containing 4μg/ml hygromycin B (Millipore Sigma) at 37°C in 5% CO2 and passaged every 3 to 4 days. UMSCC47 and UMSCC104 cell lines were obtained from Millipore Sigma (Catalog # SCC071 and # SCC072 respectively). UMSCC47 cells were grown in Dulbecco’s modified Eagle’s medium (DMEM) (Invitrogen) supplemented with 10% (vol/vol) fetal bovine serum (FBS) (Invitrogen). UMSCC104 cells were grown in Eagle’s minimum essential medium (EMEM) (Invitrogen) supplemented with nonessential amino acids (NEAA) (Gibco) and 10% (vol/vol) FBS. All cells were routinely checked for mycoplasma contamination. For protein and RNA analyses, 1 × 10^6^ cells were plated onto 100-mm2 plates, trypsinized and washed twice with 1X phosphate-buffered saline (1X PBS).

### SYBR green qRT-PCR

RNA was isolated using the SV Total RNA isolation system (Promega) following the manufacturer’s instructions. 2 μg of RNA was reverse transcribed into cDNA using the high-capacity reverse transcription kit (Applied Biosystems). cDNA and relevant primers were added to PowerUp SYBR green master mix (Applied Biosystems) and real-time PCR was performed using 7500 Fast real-time PCR system. The primer sequences utilized were as follows: Twist1 Forward (F), 5’-GTCCGCAGTCTTACGAGGAG-3’; Twist1 Reverse (R), 5’-GCTTGAGGGTCTGAATCTTGCT-3’, HPV16 E2 F, 5’-ATGGAGACTCTTTGCCAACG-3’; HPV16 E2 R, 5’-TCATATAGACATAAATCCAG-3’, VIM (Vimentin) F, 5’-GACGCCATCAACACCGAGTT-3’; VIM R, 5’-CTTTGTCGTTGGTTAGCTGGT-3’, CDH2 (N-Cadherin) F, 5’-AGCCAACCTTAACTGAGGAGT-3’; CDH2 R, 5’-GGCAAGTTGATTGGAGGGATG-3’.

### Decitabine treatment

N/Tert-1 cells were plated at a density of 1.5 × 10^5^ in 6-well plates (60-mm2/well). The following day, cells were treated with 1μM Decitabine or 1μM DMSO for 72-hours as previously described (34). Afterwards, the cells were harvested and processed for qRT-PCR as described above.

### Immunoblotting

N/Tert-1, UMSCC47 and UMSCC104 cells were trypsinized, washed with 1X PBS and respuspended in 2x pellet volume protein lysis buffer (0.5% Nonidet P-40, 50mM Tris [pH 7.8], 150 mM NaCl) supplemented with protease inhibitor (Roche Molecular Biochemicals) and phosphatase inhibitor cocktail (Sigma). Cell suspension was lysed for 30 min on ice and then centrifuged for 20 min at 184,000 rcf at 4°C. Protein concentration was determined using the Bio-Rad protein estimation assay. 25μg protein samples were boiled in equal volume 2x Laemmli sample buffer (Bio-Rad). Samples were run down a Novex 4-12% Tris-glycine gel (Invitrogen) and transferred onto a nitrocellulose membrane (Bio-Rad) at 30V overnight using the wet blot method. Membranes were blocked with Odyssey (PBS) blocking buffer (diluted 1:1 with 1X PBS) at room temperature for 1 hour and probed with indicated primary antibody diluted in Odyssey blocking buffer. Membranes were then washed with PBS supplemented with 0.1% Tween (PBS-Tween) and probed with the indicated Odyssey secondary antibody (goat anti-mouse IRdye 800CW or goat anti-rabbit IRdye 680CW) diluted in Odyssey blocking buffer at 1:10,000. Membranes were washed and underwent infrared scanning using the Odyssey CLx Li-Cor imaging system. Immunoblots were quantified using ImageJ. The following primary antibodies were used for immunoblotting: HPV16 E2 (TVG 261) at 1:1000 dilution from Abcam; Glyceraldehyde-3-phosphate dehydrogenase (GAPDH) (sc-47724) and p53 (sc-47698) at 1:1000 dilution from Santa Cruz Biotechnology and Twist1 (25465-1-AP) at 1:300 from Proteintech.

### Chromatin Immunoprecipitation (ChIP)

N/Tert-1 cells were plated at a density of 2 μ 10^6^ in 150mm2 plates. The following day, the cells were harvested via scraping and processed for chromatin as previously described (72). Chromatin concentration was determined with a NanoDrop spectrophotometer. Approximately 100μg of chromatin was used per antibody experiment. The following antibodies were used for ChIP: 2μl sheep anti-HPV16 E2 (amino acids 1 to 201) prepared and purified by Dundee Cell Products, United Kingdom; 2μg rabbit anti-Histone H3K9me2 (Abcam, ab1220). Chromatin was then processed for qPCR. ChIP DNA primers: Twist1 F, 5’-TCAGGCCAATGACACTGCT-3’; Twist1 R, 5’-GACGGTGTGGATGGCCCCGA-3’.

### Transcription Assay

5 × 10^5^ N/Tert-1 cells were plated out on 100-mm2 plates and transfected 24 hours later with either 1μg HPV16 E2 plasmid and 1μg pTWIST1-Luc, or 1μg pTWIST1-Luc alone using Lipofectamine 3000 (ThermoFisher Scientific) according to the manufacturer’s instructions as previously described (73). pTWIST1-luc contains the human promoter and has been described previously (74, 75). Briefly, the cells were harvested 72-hours post transfection utilizing the Promega reporter lysis buffer and analyzed for luciferase using the Promega luciferase assay system. Concentrations were normalized to protein levels, as measured by the Bio-Rad protein estimation assay mentioned above. Relative fluorescence units were measured using the BioTek Synergy H1 hybrid reader.

### Wound healing assay

2.5 × 10^5^ N/Tert-1 cells were plated at the center of 6-well plates (60-mm2/well). The cells were left to grow to confluency. Afterwards the monolayer was scratched using a 1000μL pipette tip, creating an ∼1mm wound. Wounds were imaged at 0, 12 and 20-hour intervals (Zeiss, Axiovert 200 M microscope). Multiple images were taken randomly along the wounds and measurements were taken from leading cell edge. Wound was measured using ImageJ software and wound healing was calculated as a ratio of the 0h timepoint.

### Statistical Analyses

The standard error was calculated from no less than three independent experiments. The only exception was in the TWIST1-Luc Transcription assay where statistical power was achieved after 2 replicates. Significance was determined using two-tailed student’s *t* test. Bonferroni correction for significance was utilized when indicated.

## Acknowledgements

This work was supported by VCU Philips Institute for Oral Health Research and the National Cancer Institute Designated Massey Cancer Center grant P30 CA016059 (IMM), and by NCI 5R01CA210911 (AA) and 5R01CA240484 (AA). We thank Dr. Lu-Hai Wang for pTWIST1-luc.

